# Surprisal contributes little beyond contextual embeddings in high-gamma ECoG encoding

**DOI:** 10.64898/2026.07.10.737662

**Authors:** Takayuki Sakuma

## Abstract

Surprisal and contextual embeddings are both derived from large language models and are widely used to predict neural responses during language comprehension, but it is unclear whether surprisal adds information beyond embeddings. We test this directly: does word surprisal improve out-of-sample prediction of high-gamma ECoG responses after GPT-2 XL contextual embeddings are included? Using public ECoG recordings from natural speech, we fit word-aligned ridge encoding models with baseline stimulus features, GPT-2 XL embeddings, and GPT-2 XL surprisal. Adding one surprisal predictor left held-out correlation essentially unchanged at the primary response window, and the increment stayed within an equivalence margin across alternative lags, residualized surprisal, and frequency-band targets; it did not uniformly meet the same criterion under banded ridge regularization or within labeled temporal subsets. This small increment suggests that surprisal does not act as an independent predictor. It is better understood as a compressed readout of the same broader predictive state that the embeddings already capture.

## 1 Introduction

Surprisal is widely used to quantify word predictability and is typically defined as the negative log probability assigned to an observed word given its preceding context [1]. In sentence-processing theory, Hale and Levy independently link this quantity to word-by-word processing difficulty [2, 3]. Smith and Levy [4] show that reading time is approximately linear in surprisal. Weissbart et al. show that word surprisal is reflected in cortical responses during continuous speech comprehension [5], while Heilbron et al. show that linguistic predictions at multiple levels are reflected in brain responses [30].

Modern language models—for example GPT-2, an autoregressive Transformer-based language model [10, 9]—assign probabilities to observed words and also generate contextual embeddings: hidden-state representations of words as processed in context. Context-sensitive language representations improve fMRI encoding models [31]: regression models that predict neural responses from stimulus features, the reverse of a decoding model, which infers stimulus properties from neural activity. Brain-based analyses have also used neural recordings to compare neural language-model representations across architectures, layers, context lengths, and attention mechanisms [32]. More broadly, integrative modeling shows that neural-network language models predict behavioral and neural responses across datasets, with next-word prediction performance closely related to brain predictivity [33]. Studies using spoken narratives bring these representations closer to natural language comprehension: GPT-2 representations map onto brain responses and predict semantic comprehension [12]. Predictive-coding work also shows that linguistic predictions are organized hierarchically over multiple levels and timescales [13]. Together, these studies establish contextual language-model representations as important predictors of neural responses during language comprehension. In these encoding models, the language model processes only the text stimulus; its outputs (embeddings and surprisal) are then used as predictors in a separate regression against the independently recorded neural response.

The most directly relevant ECoG evidence comes from natural narrative studies. Goldstein et al. show that the human brain and autoregressive language models share principles of next-word prediction, post-onset surprise, and contextual word representation during natural narrative comprehension [11]. Subsequent work shows that GPT-2 contextual embeddings share geometric structure with inferior frontal gyrus (IFG) brain embeddings [34], and that the layer-wise hierarchy of large language models aligns with the temporal dynamics of cortical responses during natural language comprehension [35]. These studies characterize the structure and timing of embedding–brain correspondence. The incremental question we test here is narrower: after GPT-2 XL embeddings are already in the model, does the same model’s word-level surprisal score improve held-out ECoG prediction? Answering this requires embeddings and surprisal in the same regression model. A simpler analysis would test surprisal alone, without embeddings in the model. Such an analysis cannot tell whether surprisal predicts neural responses on its own, or only because it correlates with the richer contextual state that embeddings capture.

A closely related preprint by de Varda et al. [37] compares surprisal, log word frequency, and word length (SFL) with contextual embeddings across eight behavioral and five neural datasets. They find that embeddings add substantial predictive information beyond SFL in neural responses, whereas SFL is generally more predictive of reading behavior. Their neural analyses model responses averaged across recording locations and participants. We instead add surprisal alone to a common baseline-plus-embedding model, fit separate ECoG electrodes, and summarize held-out prediction increments across participants. The studies therefore address related predictive questions with different features, neural measures, and aggregation procedures.

We address this incremental question with public electrocorticography (ECoG) recordings from natural speech. ECoG measures intracranial cortical activity. High-gamma activity (power in roughly the 70–200 Hz range) is often used as a local index of cortical engagement, thought to reflect nearby neuronal population firing rather than large-scale oscillatory synchrony. The open “Podcast” dataset provides ECoG recordings from participants listening to a natural spoken narrative, with stimulus features and neural recordings aligned to the same transcript [6]. Because the embedding vector already contains lexical, syntactic, semantic, and contextual information, we expect the scalar surprisal score to behave as a compressed readout of that same GPT-2 XL processing stream. We therefore use a direct incremental-prediction test: does adding GPT-2 XL surprisal to baseline features and GPT-2 XL embeddings improve held-out prediction of word-aligned neural responses?

The analysis separates marginal association from incremental predictive value. In the primary shared-ridge model with nine participants, adding surprisal changes the participant-average held-out correlation by Δ*r* = 0.000286, with its 90% confidence interval inside the ± 0.001 margin. Pointwise equivalence also holds across fixed and training-selected lags, after residualizing surprisal, for alternative scalar summaries, and across high-gamma, theta, and beta targets. Feature-block-specific regularization and labeled regional subsets yield wider or larger estimates that do not uniformly meet the same criterion, while controlled spike-in analyses show that the procedure can recover added surprisal-shaped signals.

## 2 Analysis and results

### 2.1 Dataset, neural response, and notation

The unit of analysis is a word event. The source data come from the open “Podcast” ECoG dataset (OpenNeuro ds005574) [6]. Nine participants listened to the same 30-minute spoken narrative while intracranial ECoG activity was recorded. The dataset is organized in BIDS/iEEG-BIDS format, a standard folder and metadata structure for neuroimaging and intracranial electrophysiology data [16, 17]. We use MNE-Python to read the released continuous recordings [18].

Channels are retained when labeled ECoG and not marked bad in either the recording or its channel metadata. The resulting sample contains 1,268 electrodes, ranging from 72 to 235 per participant. The overall analysis includes all retained recording sites rather than selecting electrodes by prediction accuracy. Regional summaries use name- and coordinate-matched anatomical labels described below.

Feature matrices are aligned to the token transcript, aggregated to words by word ID, and merged after verifying the word strings. The common sample contains 5,028 of the 5,136 transcript words; 108 words lack phonetic features. All models in the main comparisons use the same sample. The GPT-2 transcript provides the word-onset times used to extract neural responses.

For each electrode, the primary response is the mean of the released 512-Hz high-gamma signal over [− 100, 100) ms relative to word onset. Two fixed secondary windows are [150, 350) and [400, 600) ms. A full-lag analysis evaluates 161 centers from − 2, 000 to +2, 000 ms in 25-ms steps, using a 200-ms window at each center. That analysis uses 5,022 common words because six edge events do not support every lag.

The baseline model combines four released feature families: mel-spectral power, phonetic features, syntactic features, and static word embeddings [6]. Their released dimensions are 160, 39, 96, and 300. Briefly, mel-spectral power is the acoustic envelope across mel-frequency bands, phonetic features are indicators for speech sounds such as consonants and vowels, and syntactic features are part-of-speech and phrase-structure indicators. The contextual representation is the released 1,600-dimensional final-layer GPT-2 XL vector, averaged across subword tokens within each word. We use *contextual embedding* to mean the transformer’s hidden-state representation of a word in its preceding context [9, 10]. Its 1,600 coordinates are used jointly as a numerical feature vector.

For notation, let *w*_*t*_ denote the observed word at position *t* and *w*_*<t*_ the preceding context. Let **e**_*t*_ *∈* ℝ^1600^ denote this contextual vector aligned to word position *t*, and let *S*_*t*_ denote word-level surprisal. Let *H*_*t*_ denote the released predictive-entropy feature after word-level aggregation; it is a scalar summary of uncertainty in the GPT-2 XL next-token distribution, with larger values indicating a more diffuse distribution over candidate next tokens. Let *Y ∈* ℝ^*T ×E*^ denote the word-by-electrode neural response matrix at a fixed response window, where *T* is the number of word events and *E* is the number of electrodes for that participant.

Following contemporary surprisal analyses [1], word-level surprisal is computed from the GPT-2 XL token transcript. GPT-2 XL tokenizes text into subword units rather than whole words, so a single word can correspond to one or more tokens. The workflow reads the token-level true_prob column, computes token-level surprisal with the natural logarithm, and aggregates tokens to words by summing surprisal values within each stimulus word. Thus, for word *w*_*t*_ represented by *J*_*t*_ GPT-2 tokens 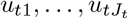, the feature used in the main analysis is

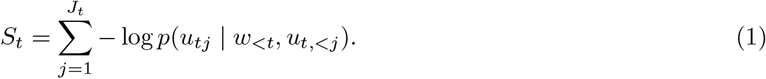

### 2.2 Contextual embeddings and surprisal

We use ridge regression [19] in a linear encoding-model framework [7, 8]. Models are fit separately for each electrode and response window. For a fixed electrode and window,

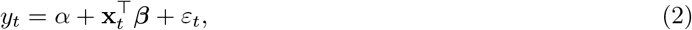

where *y*_*t*_ is the neural response to word *t* for that electrode–window pair, and **x**_*t*_ contains the predictors included in a given model specification (the exact feature composition of each named model, including *M*_base_, is listed in Table 1). The main comparison is

**Table 1:** Primary model comparison for high-gamma responses across all electrodes (nine participants). Values are participant-weighted means of held-out Pearson correlations at the 0-ms-centered response window. The reference for Δ*r* is *M*_emb_.

| Model | Feature set and interpretation | Mean *r* | Δ*r* vs. *M*<sub>emb</sub> |
|---|---|---|---|
| *M*<sub>base</sub> | Baseline stimulus features **b**<sub>*t*</sub> only: mel-spectral, phonetic, syntactic, and static word-embedding features. | 0.030438 | — |
| *M*<sub>emb</sub> | *M*<sub>base</sub> plus the GPT-2 XL contextual embedding **e**<sub>*t*</sub> for each word. | 0.038465 | Reference |
| *M*<sub>emb+surp</sub> | *M*<sub>emb</sub> plus scalar word-level surprisal *S*<sub>*t*</sub>. | 0.038751 | +0.000286 |
| *M*<sub>emb+rot</sub> | *M*<sub>emb</sub> plus 1 − cos(**e**<sub>*t*</sub>, **e**<sub>*t*−1</sub>). | 0.038375 | -0.000089 |
| *M*<sub>emb+step</sub> | *M*<sub>emb</sub> plus ‖**e**<sub>*t*</sub> − **e**<sub>*t*−1</sub>‖<sub>2</sub>. | 0.038371 | -0.000094 |

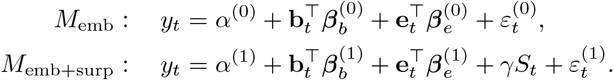

Here **b**_*t*_ and **e**_*t*_ denote the baseline and GPT-2 XL contextual-embedding features for word *t* after the training-fold preprocessing steps (standardization and, where applicable, PCA dimensionality reduction), and *S*_*t*_ is the scalar surprisal feature defined in Equation (1). Adding surprisal therefore means adding one standardized predictor column to the design matrix, with coefficient *γ*. It does not require expanding *S*_*t*_ to match the dimensionality of the embedding space. Below, *M*_base_ denotes the corresponding baseline-only model using **b**_*t*_ without **e**_*t*_ or *S*_*t*_.

Out-of-sample performance is evaluated with contiguous five-fold blocked cross-validation. A 25-word gap and a 0.25-second onset-time guard separate training and held-out blocks. Five temporally ordered inner folds apply the same exclusion rules and select the ridge penalty separately for each model, electrode, and response window. Standardization and principal component analysis (PCA) are fit within each inner training set and refit on the complete outer training set. PCA retains 50 dimensions for the spectral, syntactic, and static embedding blocks, 30 for phonetic features, and 100 for contextual embeddings. Scalar features are standardized without PCA. Penalties are selected from {1, 10, 10^2^, 10^3^, 10^4^, 10^5^, 10^6^ }.

For a fitted model *M*, let *r*(*M*) denote the held-out Pearson correlation for one electrode and outer fold. We average across folds, then electrodes within participant, and finally participants with equal weight. The main contrast is

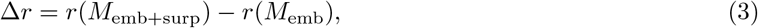

where *r*(·) is the held-out Pearson correlation defined above. Equation (2) defines the regression model fitted for a given feature set; Equation (3) compares the held-out prediction accuracy of the two fitted models. Tables and figures use *M*_base_, *M*_emb_, *M*_emb+surp_, and Δ*r* with these meanings unless otherwise stated. A positive Δ*r* means that surprisal improves prediction beyond contextual embeddings.

For the present analysis, we define practical equivalence as a participant-average change in held-out correlation within ± 0.001, a magnitude we treat as negligible on the held-out correlation scale, and assess this criterion using the two one-sided tests (TOST) procedure [20]. Statistical inference is conducted across participants, not electrodes. We report point estimates, 90% confidence intervals for the equivalence tests, and 95% *t* confidence intervals across participants.

Across all retained electrodes, *M*_base_ achieves mean held-out correlation *r* = 0.030438, while *M*_emb_ increases it to *r* = 0.038465 (Figure 1; Table 1). Adding surprisal yields *r* = 0.038751. Rotation and step size do not improve the participant average.

**Figure 1:**
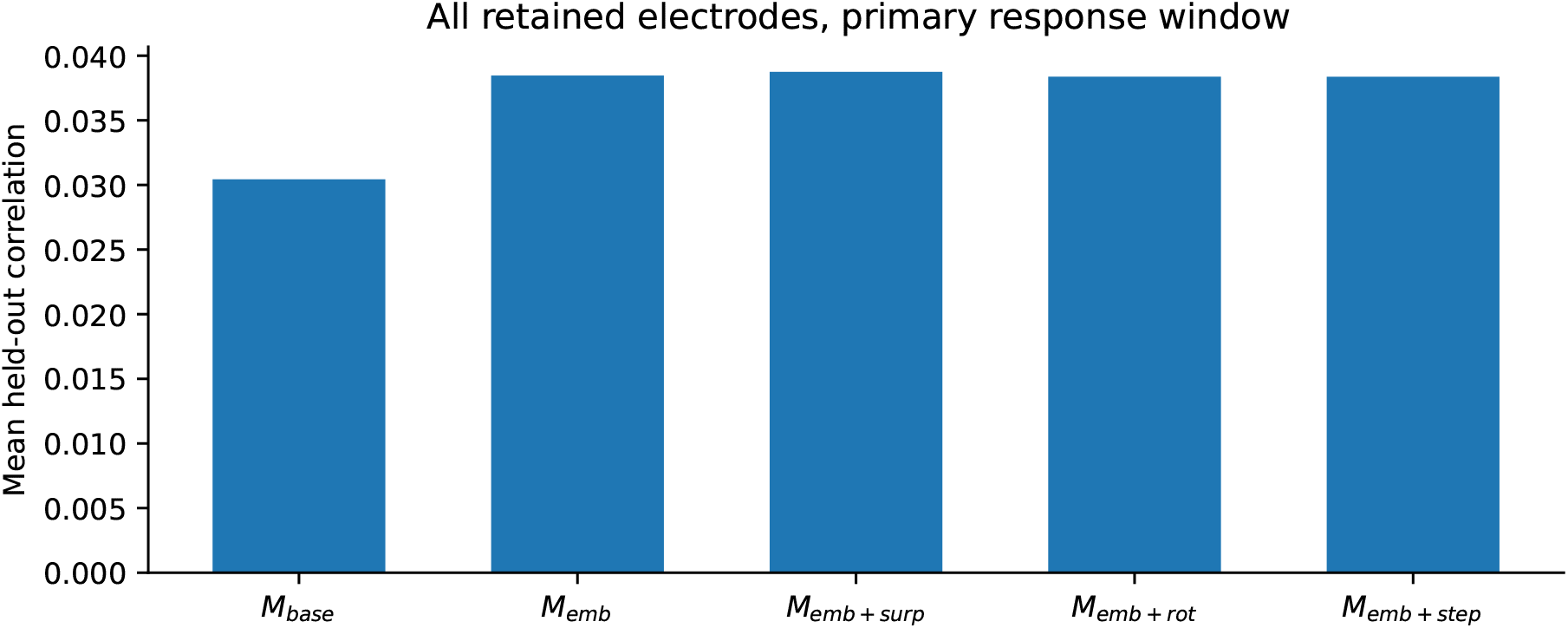
Main model comparison across all electrodes. Bars show participant-weighted mean held-out Pearson correlations at the primary response window. Exact values and model definitions are reported in Table 1.

**Figure 2:**
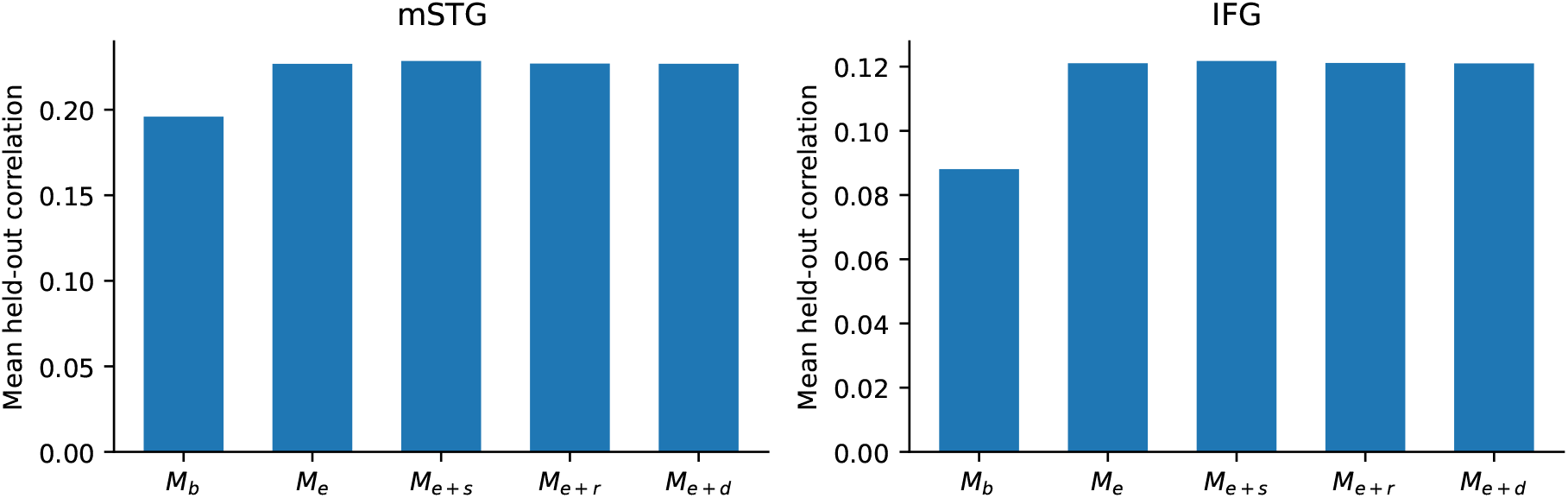
Region-specific model comparisons. Bars show held-out correlations averaged within participant and then across participants represented in the matched mSTG and IFG subsets. *M*_*b*_, *M*_*e*_, *M*_*e*+*s*_, *M*_*e*+*r*_, and *M*_*e*+*d*_ denote the baseline, embedding, surprisal, rotation, and step-size models in Table 1.

Table 1 uses the same notation as the regression equations. *M*_base_ contains the non-contextual baseline features, *M*_emb_ adds the GPT-2 XL contextual embedding, and *M*_emb+surp_ is the main surprisal-augmented model. The last two rows are secondary models that add scalar summaries of how the contextual embedding changes from one word to the next.

The same models are summarized in matched anatomical subsets. The mSTG subset contains 24 electrodes from five participants, and the IFG subset contains 55 electrodes from six participants. Superior temporal gyrus and inferior frontal gyrus are used as regions of interest because they are canonically implicated in auditory speech processing and language comprehension, respectively. At the primary window, adding surprisal changes the mSTG mean from 0.226829 to 0.228486 and the IFG mean from 0.121057 to 0.121756. Across the three fixed windows, mSTG increments range from 0.0014 to 0.0031, while IFG increments range from 0.0004 to 0.0007. Neither regional subset satisfies the ± 0.001 equivalence criterion at the primary window: the 90% interval is [0.000701, 0.002613] for mSTG and [0.000210, 0.001189] for IFG. None of the three mSTG windows meets the criterion; for IFG, the 0- and 250-ms windows do not meet it, whereas the 500-ms window does. The all-electrode equivalence result therefore does not extend to these labeled subsets. The regional differences themselves are not tested, and the source labels do not constitute a complete parcellation or an independent language localizer.

Adding *S*_*t*_ to *M*_emb_ changes the all-electrode participant mean by 0.000286 (95% CI: [− 0.000045, 0.000617]). The 90% confidence interval, [0.000019, 0.000553], lies entirely within the equivalence margin of ± 0.001, and the TOST procedure supports practical equivalence to zero (*p* = 5.4 *×* 10^−4^). In this word-aligned encoding model, scalar word surprisal therefore adds at most a small amount of out-of-sample predictive information beyond contextual embeddings. Seven of nine participant increments are positive and two are negative (Figure 3). Individual means range from −0.000095 to 0.001241.

**Figure 3:**
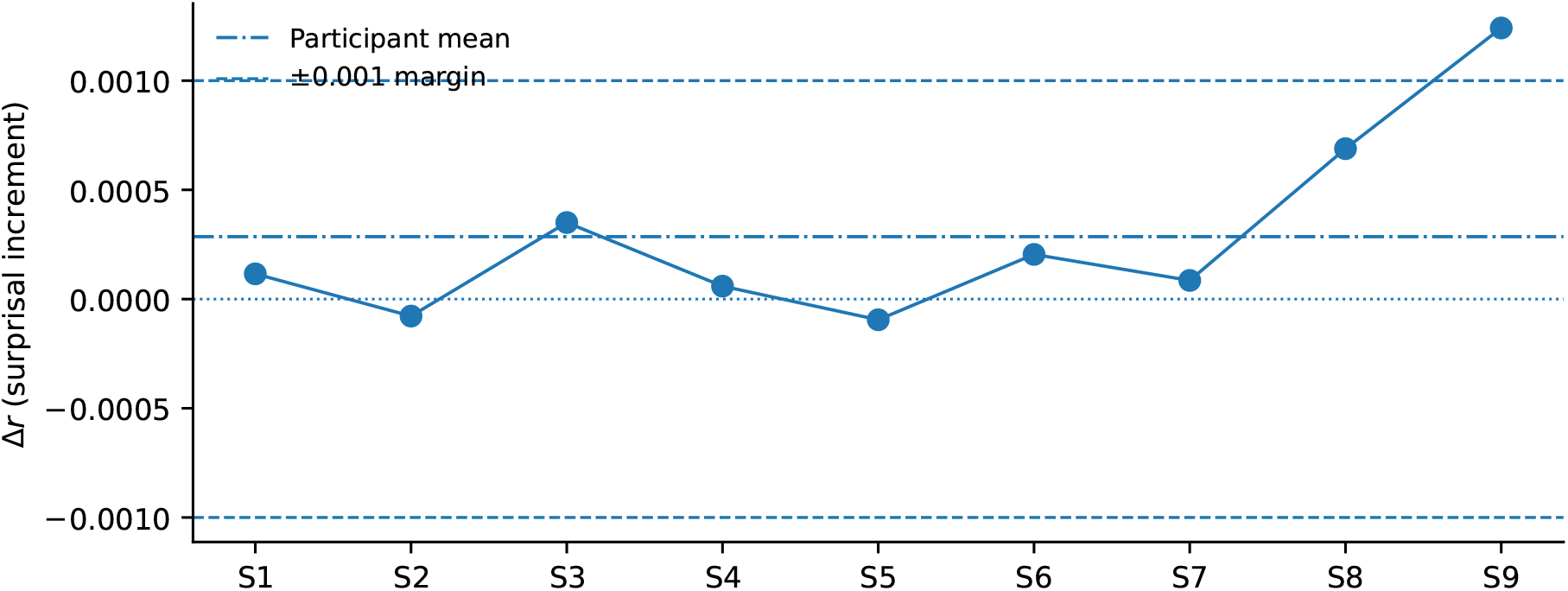
Participant-level incremental effect of adding surprisal. Each point shows Δ*r* for one participant at the primary response window. The dash-dotted line is the participant mean, dashed lines mark the equivalence margin of ±0.001, and the dotted line marks zero.

### 2.3 Lag and feature-block analyses for the surprisal contrast

The full-lag analysis repeats Equation (3) at 161 window centers. Because all candidate windows must be available for each event, it uses the 5,022-word common sample described above rather than the 5,028-word primary sample. Its fixed 0-ms estimate is 0.000276, close to but not numerically identical to the headline estimate of 0.000286. It uses a 4.2-second onset guard, which spans the candidate lag range plus window width and prevents direct overlap between training and test response windows across candidate lags. Mean Δ*r* ranges from − 0.000190 to +0.000281 over the full grid. At every window center, the participant-level 90% interval lies within the ± 0.001 margin; a Bonferroni-adjusted simultaneous 95% band exceeds the margin at 10 of 161 centers, so the pointwise equivalence should not be read as a simultaneous guarantee.

Table 2 also reports three peak-lag rules. For each participant and outer fold, the lag is selected using only inner-validation performance for the contextual-embedding model, the spectral-feature model, or surprisal alone. The last rule gives the surprisal feature its most favorable timing before conditioning on embeddings. The predictors are unchanged; only the response window differs across rows. The conditional *M*_emb+surp_ − *M*_emb_ contrast is then evaluated on the outer held-out words. Mean increments remain small under all three rules (0.000039, 0.000275, and 0.000248), and each 90% interval lies within the ±0.001 margin.

**Table 2:**
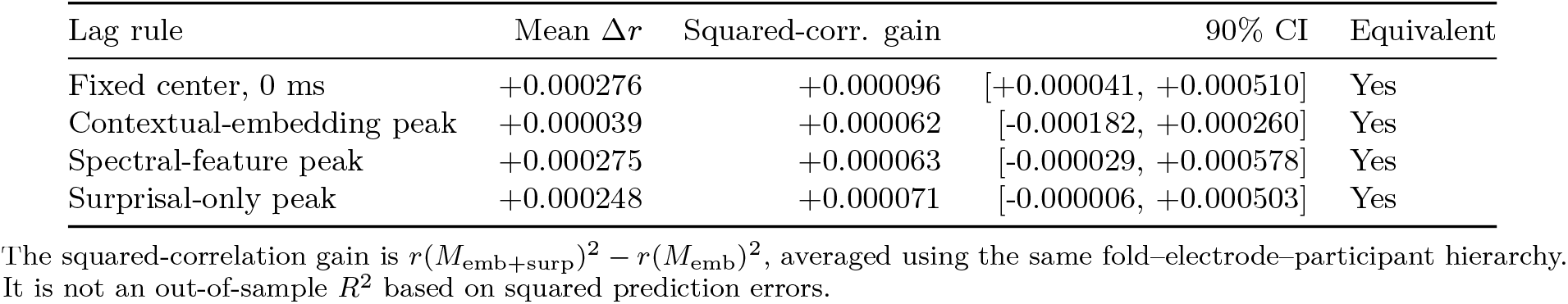
Lag-resolved analyses of the main surprisal contrast (nine participants). The fixed center and the three peak rules use the 5,022-event full-grid sample. Peaks are selected within each outer training set using inner validation only. Equivalence is assessed against ±0.001.

Table 3 asks how much feature-specific information remains after conditioning on the other block. Adding the contextual embedding to a baseline-plus-surprisal model increases mean correlation by 0.007500; its 90% interval, [0.000401, 0.014598], is not contained in the equivalence margin. In the reverse direction, surprisal increases mean correlation by 0.000286 and meets the equivalence criterion. Residualizing surprisal with respect to the current-word embedding within each outer training set yields a mean increment of 0.000305 (90% CI: [0.000006, 0.000605]). A previous-word residualization check gives 0.000213 (90% CI: [0.000009, 0.000417]). Both residualized-surprisal intervals lie within the ± 0.001 margin. Thus the asymmetry remains: the embedding adds information not captured by surprisal, while surprisal adds little after the embedding is included.

**Table 3:**
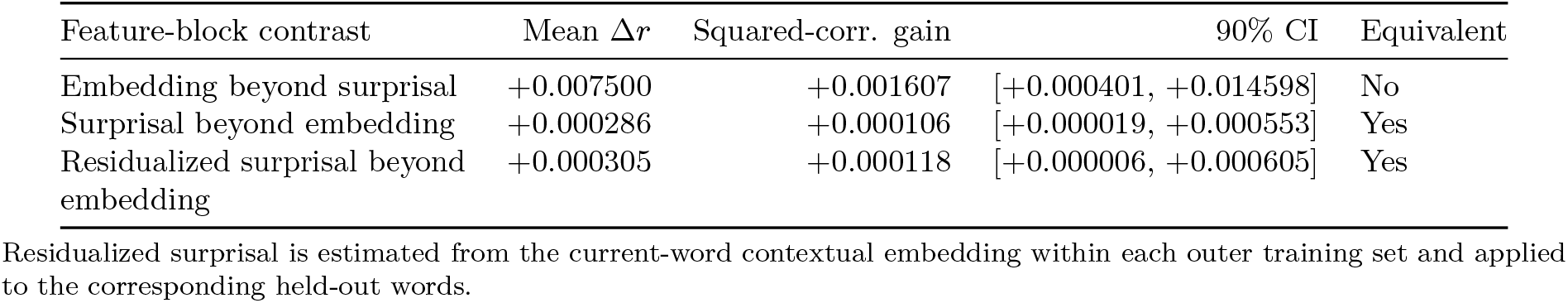
Feature-block overlap contrasts at the primary response window (nine participants). Each row compares a larger model with the indicated smaller reference model. Equivalence is assessed against±0.001.

### 2.4 Exploratory contextual-update summaries

The main analysis concerns scalar surprisal. To test whether the small increment is specific to probability-derived surprisal, we add three one-dimensional summaries of GPT-2 XL state change to *M*_emb_: embedding step size, embedding rotation, and entropy drop. For each word, we compute the embedding update

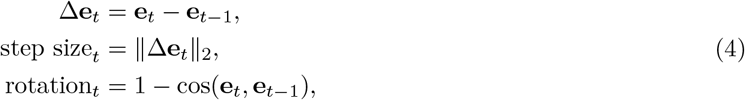

and entropy drop is max(*H*_*t* −1_ − *H*_*t*_, 0). The updates are computed in the original narrative order before the analysis intersection is formed. Each scalar update measure is added separately to *M*_emb_, and the change in held-out prediction is evaluated using the same Δ*r* contrast as in Equation (3).

The analysis does not show a positive participant-average increment for rotation or step size. Their mean changes are − 0.000089 and − 0.000094, and entropy drop changes the mean by +0.000036 (Table 4; Figure 4). The 90% intervals for all three measures lie within the ± 0.001 margin. These results do not support the claim that the simple update summaries are more predictive than surprisal.

**Table 4:** Incremental performance of candidate contextual-state-update measures. Each scalar feature is appended separately to *M*_emb_ at the primary response window. Equivalence is assessed against ±0.001.

| Feature | Mean Δ*r* | Median | Positive | Negative | 90% CI | Equivalent |
|---|---|---|---|---|---|---|
| Rotation | -0.000089 | -0.000106 | 4 | 5 | [-0.000261, +0.000083] | Yes |
| Step size | -0.000094 | -0.000048 | 4 | 5 | [-0.000352, +0.000165] | Yes |
| Entropy drop | +0.000036 | +0.000003 | 5 | 4 | [-0.000069, +0.000141] | Yes |

**Figure 4:**
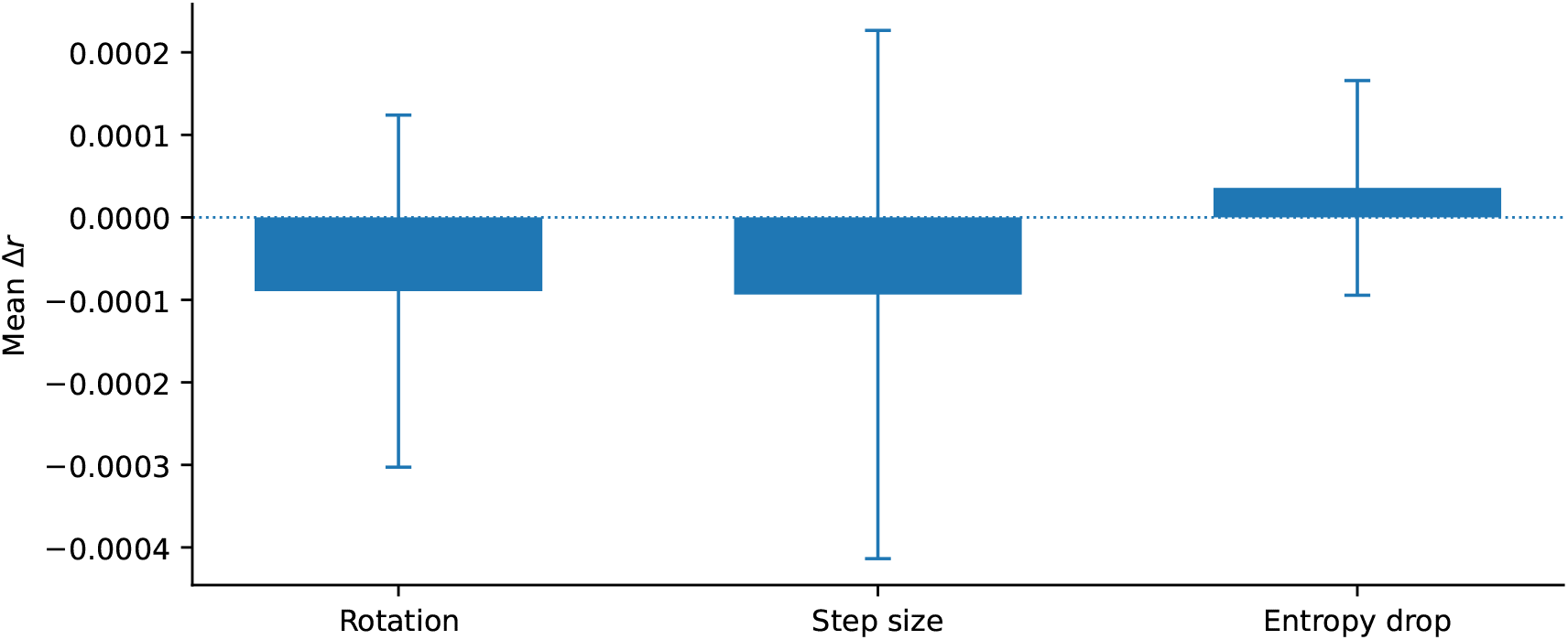
Incremental performance of candidate update measures. Bars show participant-average changes in held-out correlation relative to *M*_emb_; error bars are 95% *t* confidence intervals.

### 2.5 Sensitivity analyses

The analyses in this subsection repeat the same core contrast, *M*_emb_ versus *M*_emb+surp_. Each check changes one element of the workflow while leaving the comparison unchanged: the scalar predictor set, the neural target, the regularization scheme, or the response matrix used for calibration.

The first sensitivity analysis compares three scalar additions to *M*_emb_: surprisal, predictive entropy, and both together. The comparisons use the same nine participants and primary response window. Mean changes are small in all three cases (Table 5; Figure 5), and all three 90% intervals lie within the ±0.001 margin.

**Table 5:** Alternative scalar additions to the contextual-embedding model (nine participants). Values are changes in held-out correlation relative to *M*_emb_ at the primary response window. Equivalence is assessed against ±0.001.

| Model compared with *M*<sub>emb</sub> | Mean Δ*r* | SD | 90% CI | Equivalent |
|---|---|---|---|---|
| *M*<sub>emb</sub> + *S*<sub>*t*</sub> | +0.000286 | 0.000431 | [+0.000019, +0.000553] | Yes |
| *M*<sub>emb</sub> + *H*<sub>*t*</sub> | -0.000071 | 0.000166 | [-0.000174, +0.000031] | Yes |
| *M*<sub>emb</sub> + *S*<sub>*t*</sub> + *H*<sub>*t*</sub> | +0.000156 | 0.000525 | [-0.000170, +0.000481] | Yes |

**Figure 5:**
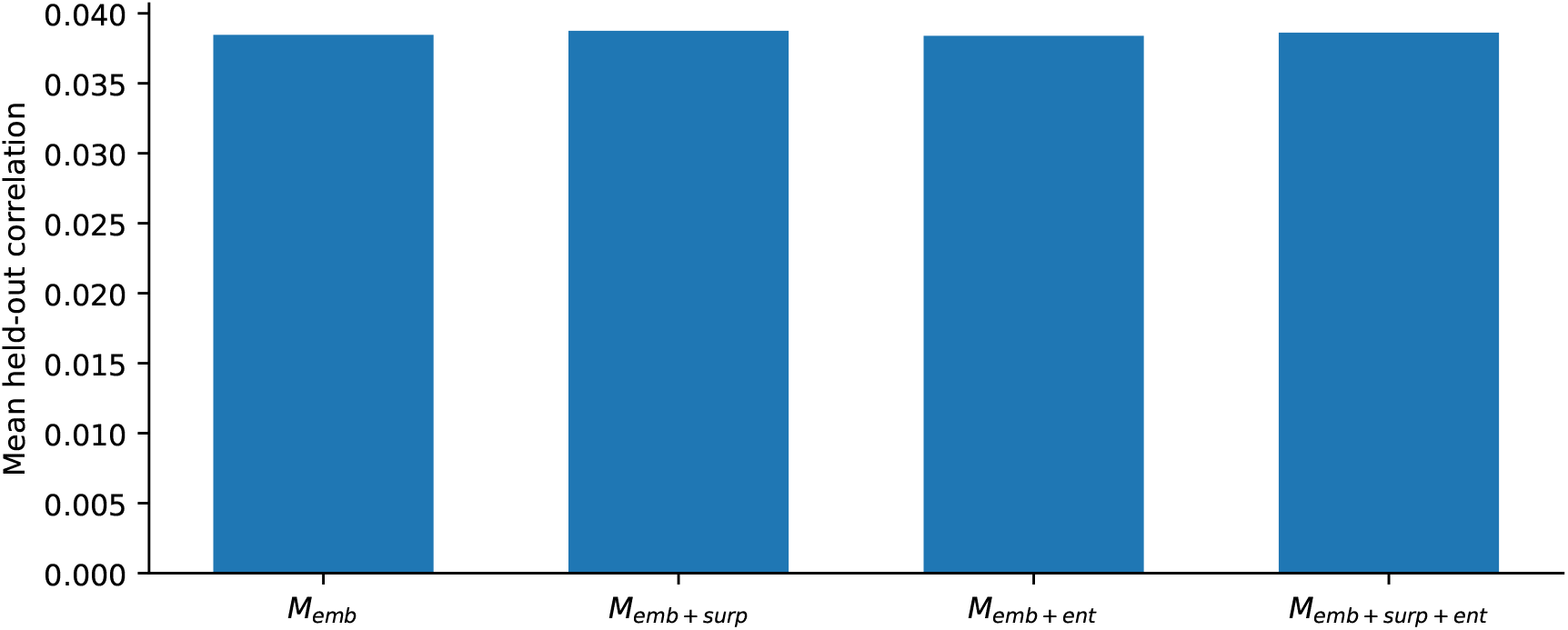
Alternative scalar additions to the contextual-embedding model. Bars show participant-weighted mean held-out correlations for *M*_emb_ and models that add surprisal, entropy, or both.

We next re-express the same ECoG channel time series as theta (4–8 Hz) and beta (13–30 Hz) Hilbert-amplitude envelopes. For each channel, the signal is band-pass filtered in the theta or beta range, and the Hilbert amplitude is used as the band-limited envelope. This applies the same filtering-and-envelope operation used for high-gamma broadband activity to lower-frequency bands. The same 5,022 words, retained electrodes, predictors, and validation design are used for each target; only the neural response changes. At the 0-ms-centered window, mean Δ*r* is 0.000276 for the matched high-gamma comparison, 0.000258 for beta, and 0.000219 for theta (Table 6). The 90% intervals for all three neural targets lie within the ± 0.001 margin. The changes are also small at the 250- and 500-ms windows.

**Table 6:** Sensitivity analysis across band-limited neural targets. The same 5,022 words, retained electrodes, predictors, and validation design are used for each target; only the neural response changes. Equivalence is assessed against ±0.001.

| Neural target | *n* | Mean Δ*r* | SD | 90% CI | Equivalent |
|---|---|---|---|---|---|
| High-gamma | 9 | +0.000276 | 0.000378 | [+0.000041, +0.000510] | Yes |
| Beta envelope | 9 | +0.000258 | 0.000382 | [+0.000020, +0.000495] | Yes |
| Theta envelope | 9 | +0.000219 | 0.000521 | [-0.000104, +0.000542] | Yes |

The main ridge models use a shared penalty for the concatenated feature vector. Because the contextual embedding is high-dimensional and surprisal is a scalar, we also use feature-space-specific regularization [21, 22]. The four baseline spaces form one penalty group, contextual embeddings another, and surprisal a third. Block weights and ridge penalties are selected in inner folds and evaluated only on outer folds.

The banded-ridge mean increment is 0.000861, with a 95% interval of [− 0.000967, 0.002690], compared with 0.000286 under shared ridge (Table 7; Figure 6). The point estimate remains small on the correlation scale, but participant variation and uncertainty are greater. Under banded ridge the 90% interval, [− 0.000613, 0.002336], extends beyond the ± 0.001 margin, and the TOST procedure does not support equivalence (*p* = 0.43). The banded-ridge analysis therefore neither confirms nor contradicts the shared-ridge equivalence result.

**Table 7:** Regularization analysis for the primary surprisal comparison. Both specifications use the same nine participants, 1,268 electrodes, 5,028 word events, and primary response window. Equivalence is assessed against ±0.001.

| Analysis | *r*(*M*<sub>emb</sub>) | *r*(*M*<sub>emb+surp</sub>) | Mean Δ*r* | SD | 90% CI | Equivalent |
|---|---|---|---|---|---|---|
| Shared ridge | 0.038465 | 0.038751 | +0.000286 | 0.000431 | [+0.000019, +0.000553] | Yes |
| Banded ridge | 0.039019 | 0.039881 | +0.000861 | 0.002378 | [-0.000613, +0.002336] | No |

**Table 8:**
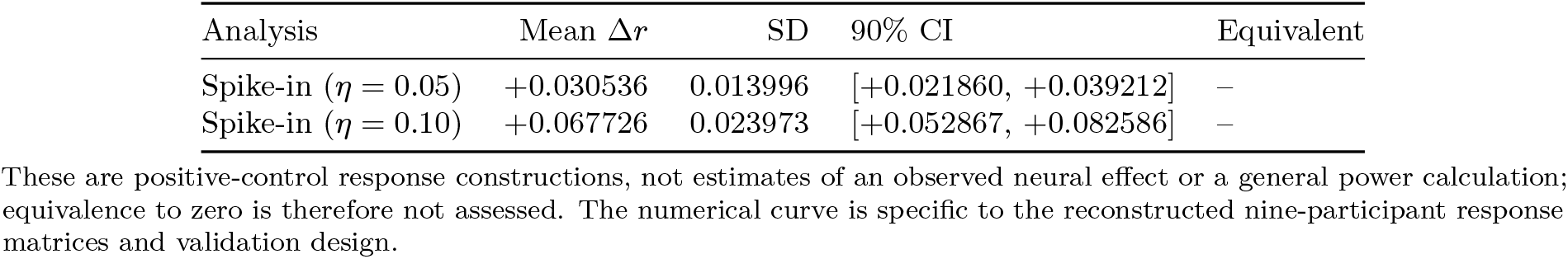
Spike-in sensitivity analysis. A controlled surprisal-shaped signal is injected into the neural response matrix and the same nested comparison is repeated.

**Figure 6:**
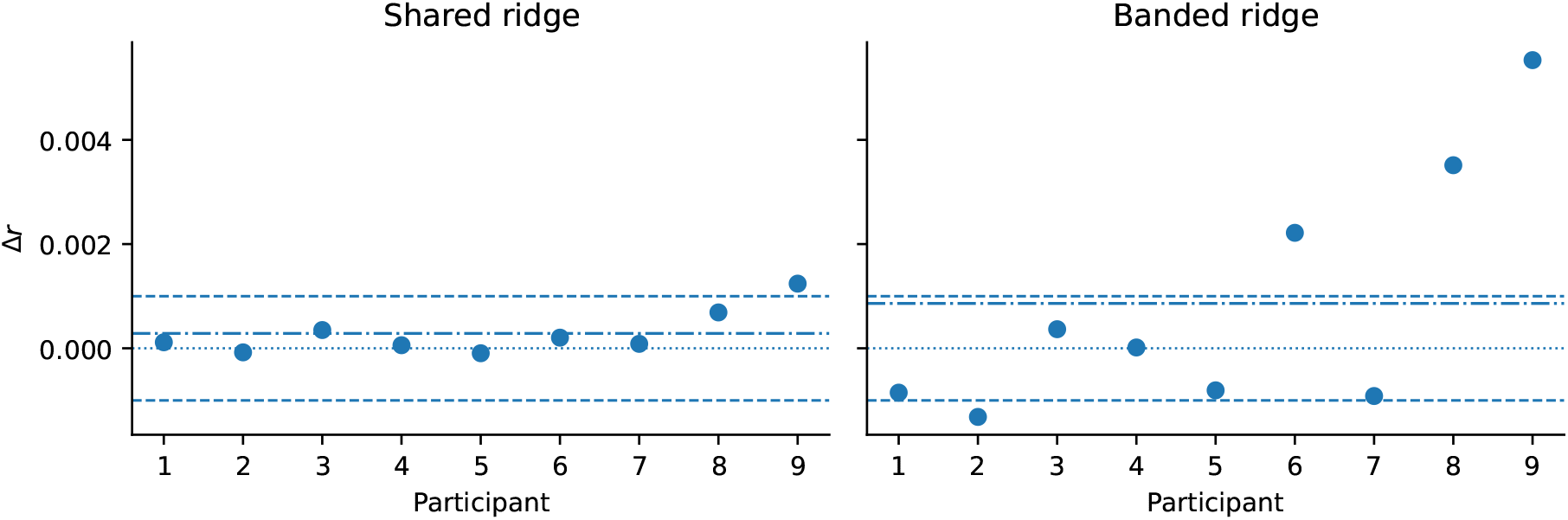
Participant-level surprisal increments under shared and banded ridge. Points are participant means across retained electrodes at the primary response window. Dash-dotted lines indicate the respective participant means, dashed lines mark the equivalence margin of ± 0.001, and dotted lines mark zero.

**Figure 7:**
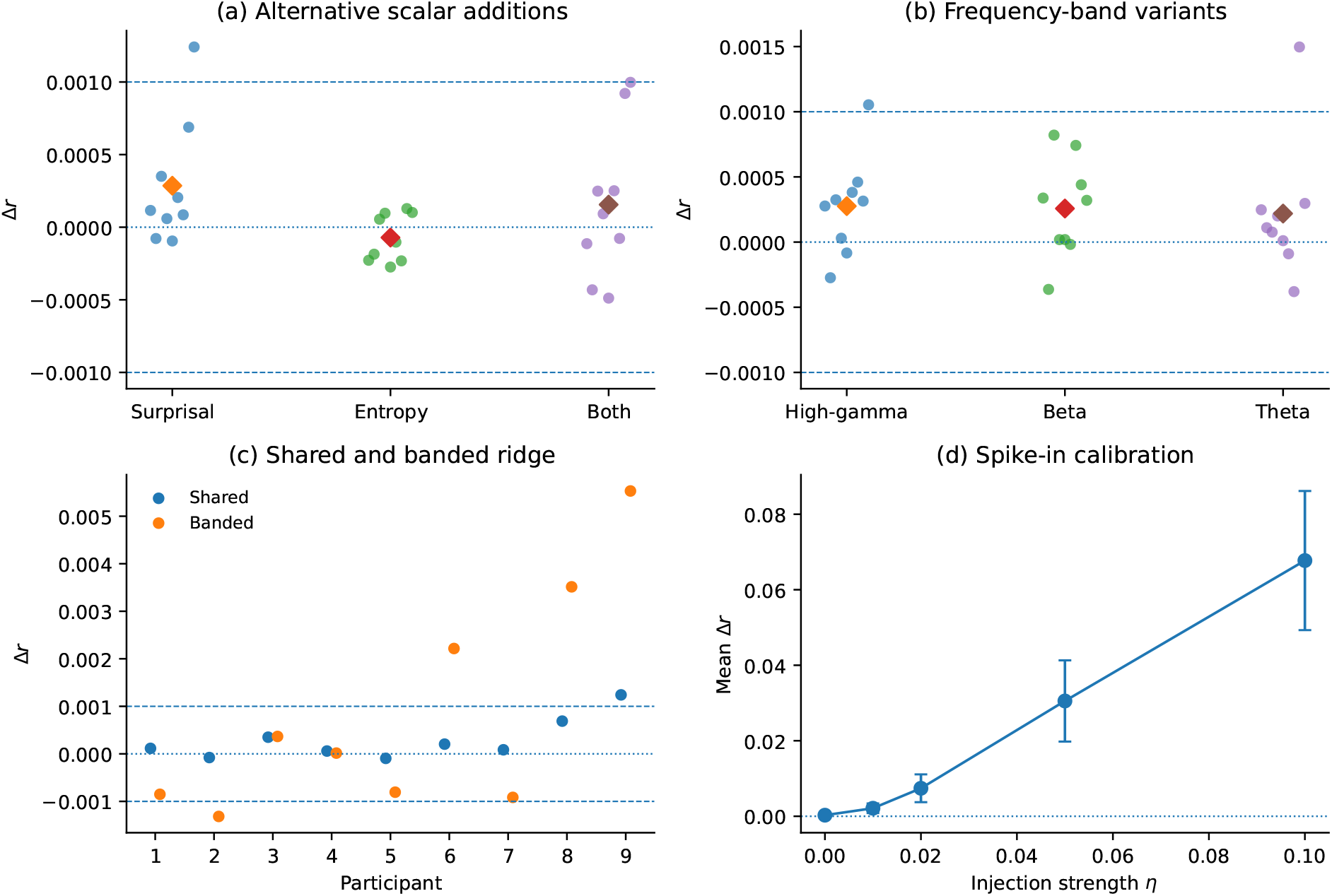
Sensitivity analyses of the main surprisal comparison. Panel (a) shows participant increments after adding surprisal, entropy, or both. Panel (b) compares the matched high-gamma, beta, and theta targets at the primary window. Panel (c) compares shared and banded ridge. Panel (d) shows the recovered increment after injecting a controlled surprisal-shaped component. Diamonds in panels (a) and (b) denote participant means; dashed lines in panels (a)–(c) mark the ±0.001 equivalence margin.

Finally, we calibrate sensitivity by adding a surprisal-shaped component to each participant’s response matrix:

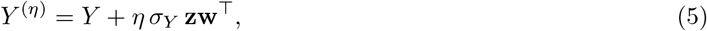

where *Y* is the original word-by-electrode response matrix, *σ*_*Y*_ is its global standard deviation, **z** is the z-scored surprisal vector across words, **w** is a fixed normalized random electrode-weight vector, and *η* (distinct from the regression coefficient *γ* in Equation (2)) takes values 0, 0.01, 0.02, 0.05, and 0.10. We retune the models for each artificial target. Mean Δ*r* rises from 0.000286 at *η* = 0 to 0.002125, 0.007413, 0.030536, and 0.067726 as *η* increases. This positive control shows that the procedure can recover the specified scalar-shaped signal; it is not a biological threshold or a general power analysis. Its numerical curve is specific to the reconstructed nine-participant response matrices and nested-validation design.

## 3 Discussion

We tested whether word surprisal improves prediction of word-aligned neural responses after contextual embeddings are already included in the model. Four findings summarize the results. First, contextual embeddings improve participant-average prediction relative to the baseline feature set. Second, adding surprisal changes the all-electrode shared-ridge mean only slightly, including across a broad lag range, after residualizing surprisal, and for theta and beta envelopes. Third, the increment is more variable under banded ridge and modestly larger in the labeled mSTG subset. Fourth, rotation, step size, entropy drop, and predictive entropy do not yield clear average improvements beyond the embedding. A controlled positive-response perturbation shows that the procedure can recover added surprisal-shaped signals.

The small overall increment matters because many surprisal findings are estimated against low-dimensional baselines or marginal associations. Those designs can show that surprisal relates to neural responses, but they do not show how much predictive value it retains beyond a contextual model of the stimulus. The present result concerns the latter question. The asymmetry in Table 3 is consistent with embeddings containing predictive information not represented by surprisal, while surprisal adds little on average after embeddings are included.

The comparison with de Varda et al. [37] is informative. Their neural results show that embeddings generally outperform the combined SFL predictors and that adding SFL to embeddings produces relatively small gains. Our result isolates surprisal within an electrode-wise ECoG analysis rather than modeling SFL jointly or averaging the neural response before fitting. The findings are qualitatively aligned, but they are not estimates of the same contrast. In particular, the regional and banded-ridge results here caution against interpreting predictive overlap as uniform across sites and specifications.

One possible explanation for the small average increment is predictive overlap between surprisal and a broader contextual representation. In autoregressive transformers, next-word probabilities are computed from contextual hidden states [9, 10]. Surprisal keeps only the probability assigned to the observed word, whereas the contextual representation preserves a high-dimensional state. Surprisal discards the probabilities assigned to every other possible next word. If neural activity tracks a distributed contextual state or a transformation of that state, then a single scalar summary has limited room to improve out-of-sample prediction. This interpretation is also compatible with predictive-coding accounts that distinguish state representations from error-related signals [14, 15], without requiring a one-to-one mapping between scalar word surprisal and neural activity. At the same time, the released contextual vector describes the word after it is processed, while surprisal comes from the distribution before that token is observed. Their common model origin therefore does not make the regression comparison mathematically redundant.

The update summaries do not provide evidence that a different scalar description of contextual change resolves this limitation. Rotation, step size, and entropy drop all have participant-average increments close to zero. The result does not rule out more informative decompositions of the embedding space, but it gives no reason to privilege these three one-dimensional summaries in the present data. Which decomposition is most informative is not settled by this analysis. Candidates include hand-constructed update measures of the kind used here, principal components, and learned decompositions such as sparse autoencoders, which have been used to recover interpretable features from language-model activations [36]. Identifying which of these best relates to neural responses is a natural target for future work.

These results narrow how scalar surprisal should be interpreted in applied settings. In the present analysis, surprisal is too compressed to stand in for the contextual or belief state relevant for neural responses during naturalistic language comprehension. Scalar surprisal remains useful as a compact reduced-form statistic, but mechanistic interpretations should distinguish it from the richer state that generates it.

The spike-in analysis addresses a procedural concern. A scalar added to the same predictor set improves held-out prediction when a component with that scalar’s shape is added to the response, and the recovered increment increases with injection strength. This establishes sensitivity to the specified construction, not to every small, sparse, or localized neural effect. Banded ridge addresses a different concern—feature groups may require different regularization—and its wider interval makes that uncertainty visible.

The averaging scheme is important. The 1,268 recording sites are retained on data-quality grounds rather than selected as language-responsive electrodes. A small all-electrode mean can coexist with larger local increments. The mSTG mean changes by 0.0014–0.0031 across the three fixed windows, but the subset includes five participants and 24 labeled electrodes, the intervals are pointwise and uncorrected, and no direct regional comparison is performed. The result therefore motivates localization rather than establishing a region-specific effect.

Canonical predictive-coding theories distinguish state representations from error representations [14, 15]. The current findings caution against mapping that distinction directly onto a scalar surprisal regressor in word-aligned high-gamma ECoG. They are not evidence against prediction-error computations in the brain. Error-related information may be distributed, multidimensional, temporally finer than word-level alignment, or expressed in signals not captured by amplitude envelopes.

The inference is bounded by the encoding model. Prior language-encoding studies address the gap between model features and neural data by using held-out prediction, regularization, temporal response modeling or lag analyses, control features, and comparisons among feature spaces [7, 11, 13, 22]. These practices reduce overfitting and define a predictive correspondence between stimulus descriptions and neural responses. Recent work on pre-onset language-encoding designs further shows that dependencies within natural language stimuli can produce prediction-like performance in passive control systems if the statistical structure of the stimulus is not handled carefully [29].

Several limitations remain. The models are linear and word-aligned rather than nonlinear or subword-resolved. All participants heard one story, so participant intervals do not establish generalization across narratives. The anatomical labels cover limited matched subsets rather than all electrodes. The frequency-band analysis uses amplitude envelopes and does not address phase or laminar structure. Future work should test whether nonlinear mappings, phase-sensitive measures, alternative language models, independent language localizers, finer temporal alignment, or richer decompositions of contextual representations reveal independent error-like signals.

More broadly, the same incremental-validity logic could extend beyond single words to narrative-level structure, such as character-state updates or event boundaries. Empirical literary studies and neurocognitive poetics emphasize that literary reading involves lexical predictability, affect, foregrounding, immersion, aesthetic response, and interpretation [23]. Naturalistic narrative studies have used literary materials such as *Harry Potter* to model reading subprocesses and affective engagement in fMRI [24, 25]. Public eye-tracking corpora such as GECO and Provo also provide behavioral outcomes for natural reading and predictability [26, 27]. Recent economic work provides a parallel example: LLMs are used to test whether completed life stories contain preference-relevant structure recoverable from human-written narratives, with validation against human responses [28]. A narrative-level follow-up could test whether LLM-derived features—character-state updates, event boundaries, affective shifts, value conflicts, and thematic changes—predict reading times, eye movements, memory, affective ratings, aesthetic response, or neural activity beyond local word surprisal. Such a study would keep the present incremental-validity design while shifting the unit of analysis from words to scenes, characters, and narrative worlds.

## 4 Conclusion

For word-aligned high-gamma responses in the shared-ridge, all-electrode analysis, contextual embeddings captured most of the predictive information available to this linear encoding model, whereas scalar surprisal contributed little beyond that state representation. This pattern held across alternative lags, feature-block redundancy checks, and frequency bands, but did not extend uniformly to banded ridge regularization or labeled regional subsets. A separate positive-control analysis confirmed that the method could recover an artificially injected surprisal-shaped component. A useful next step is to move from word-level prediction to narrative-state updating in literary comprehension, extending the same incremental-validity logic to character-state updates, event boundaries, and other narrative-level structure.

## Data and code availability

The recordings, channel metadata, transcript, and stimulus features are publicly available as OpenNeuro ds005574 [6]. The accompanying package contains the analysis code, input definitions, word-ID correspondence, participant-level results, and scripts for the reported tables and figures. The source recordings were collected by the dataset investigators; no new participant data were collected for this study.

## Declaration of AI-assisted technologies

During the preparation of this manuscript, the author used generative AI and AI-assisted tools to assist with manuscript drafting, language refinement, code drafting and debugging, and discussion of the presentation of the research. The author directed the project, reviewed, revised, and verified the manuscript and code, and takes full responsibility for the final manuscript.

